# Genotypic sex shapes maternal care in the African Pygmy mouse, *Mus minutoides*

**DOI:** 10.1101/2022.04.05.487174

**Authors:** Louise D. Heitzmann, Marie Challe, Julie Perez, Laia Castell, Evelyne Galibert, Agnes Martin, Emmanuel Valjent, Frederic Veyrunes

## Abstract

Sexually dimorphic behaviours, such as parental care, have long been thought to be driven mostly, if not exclusively, by gonadal hormones. In the past two decades, a few studies have challenged this view, highlighting the direct influence of the sex chromosome complement (XX vs XY or ZZ vs ZW). The African pygmy mouse, *Mus minutoides*, is a wild mouse species with naturally occurring XY sex reversal induced by a third, feminizing X* chromosome, leading to three female genotypes: XX, XX* and X*Y. Here, we show that sex reversal in X*Y females shapes a divergent maternal care strategy from both XX and XX* females, rather than altering care quality. In addition, we show that sex reversal may also impact the dopaminergic system in the anteroventral periventricular nucleus of the hypothalamus, consistent with one component of maternal care: pup retrieval. Combining behavioural ecology and neurobiology in a rodent subject to natural selection, we evaluate potential candidates for the neural basis of maternal behaviours and strengthen the underestimated role of the sex chromosomes in shaping sex differences in brain and behaviours. All things considered, we further highlight the emergence of a third sexual phenotype, challenging the binary view of phenotypic sexes.

## Introduction

Parental care, though not universal across the animal kingdom, is fundamental in many species for offspring’ survival and thus, reproductive success. In mammals, parental care is skewed towards females with an obligatory lactation and greater investments, making it one of the most sexually dimorphic behaviour (see Royle et al., 2012). Male care for offspring is rare (5-10% of mammalian species), mainly observed in monogamous species and always associated with biparental care (Clutton-Brock, 1991; Lukas & Clutton-Brock, 2013). On a proximate scale, this dimorphism as well as other sexually dimorphic traits, was long thought to be a product of gonadal steroid hormones only, acting to shape brain neural circuits (Arnold & Gorski, 1984; Breedlove et al., 1999). In the past two decades, several studies have challenged this view, providing increasing evidence for the direct role of the sex chromosome complement (XX or XY) on sexually dimorphic phenotypes (Arnold, 2004; Gatewood et al., 2006; Mank, 2009; Ngun et al., 2011; Skuse, 2006; Rice, 1984). More precisely, genetically modified mice have allowed to dissociate individuals’ gonadal and genotypic sexes such as in the four core genotypes (FCG) model, where the *Sry* gene has been deleted and translocated on an autosome (XX and XY^-^ females, XX*Sry* and *XY^-^Sry* males) (see Arnold, 2020). These advances have been crucial to decipher the direct role of both hormones and sex chromosomes on social behaviours (De Vries et al., 2002; Gatewood et al., 2006; Kopsida et al., 2013; Ngun et al., 2014; Skuse, 2006). However, very few studies have examined how sex chromosomes shape parental care and underlying brain neural circuits (Gatewood et al., 2006).

In rodents for instance, males tend to have more cell bodies and greater fiber densities of vasopressin expressing neurons, a neuropeptide involved in crouching over the pups or maternal aggression among others, likely to result from an imbalance in X chromosome number between the sexes (Crenshaw et al., 1992; De Vries & Miller, 1998; De Vries et al., 2002; Gatewood et al., 2006; Bosch & Neumann, 2012; Cox et al., 2015; Miller et al., 1989;Rood et al., 2008; Rood et al., 2013). Closely related to vasopressin, oxytocin is another substantial neuropeptide for parental care, well-known for its involvement in lactation, but also for enhancing pup retrieval and maternal aggression in females (Gimpl & Fahrenholz, 2001; Pedersen et al., 2006; Okabe et al., 2017; Scott et al., 2015; Bosch & Neumann, 2012). However, the impact of the sex chromosome complement on oxytocin in a parental context in the FCG or other mouse model has never been investigated. While the FCG has been a breakthrough in pointing out the value of the sex complement on parental behaviours, the common view of the system is far from complete considering the complexity of parenting behaviours as well as the various underlying neural pathways. Additionally, studies have only focused on genetically modified animals so far -which may have some limitations (e.g. absence of Sry in XY-females)-, where naturally occurring sex reversal systems subject to natural selection could bring insightful information especially in terms of brain and behaviours’ evolution.

*In natura*, a handful of rodent species display atypical sex determination systems with loss of the Y chromosome in males, or females carrying a Y chromosome (Saunders & Veyrunes, 2021). Among them, the African pygmy mouse, *Mus minutoides*, is peculiarly interesting: a third sex chromosome, called X*, has evolved from a mutation on the X chromosome, and induces sex reversal of X*Y individuals (Veyrunes et al., 2010a). Thus, there are three distinct female genotypes in populations, XX, XX* and X*Y, while all males are XY. Sex-reversed X*Y females are fully fertile (Rahmoun et al., 2014), and have a higher reproductive success than XX and XX* females in laboratory conditions (Saunders et al., 2014). In addition, when compared to XX and XX* females, X*Y females are more aggressive towards intruders, display reduced anxiety, similar to that of males, and have a greater bite force, also comparable to males (Saunders et al., 2016; Ginot et al., 2017). These male-typical behaviours observed in X*Y individuals, which are unlikely to be testosterone dependant (Veyrunes et al., 2022), suggest that the Y chromosome and/or dosage imbalance in X chromosomes masculinize some neural circuits in the brain of *M. minutoides*. The highly contrasted phenotypes between female genotypes thus represent a tremendous opportunity to understand the neural bases of sexually dimorphic behaviours. Furthermore, the dissociation of gonadal and genotypic sexes makes the African pygmy mouse a key species to investigate the effect of sex chromosomes on sex differences in brain and behaviours. In this study, we used an integrative approach to investigate the effect of genotypic sex on maternal care in *M. minutoides*, from behaviours to candidate brain neural circuits (Fig. 1). Because we show, here, that sex reversal in *M. minutoides* influences pup retrieval, maternal aggression, and nest building behaviours, we focused neural investigations on neurons expressing oxytocin and vasopressin in the paraventricular nucleus of the hypothalamus (PVN), a region and neuropeptides regulating these behaviours (Bendesky et al., 2017, Pedersen et al., 2006; Okabe et al., 2017; Scott et al., 2015; Bosch & Neumann, 2012). Since oxytocin neurons in the PVN are modulated by dopamine neurons in the anteroventral periventricular nucleus of the hypothalamus (AVPV) (Scott et al., 2015) and that dopamine levels can be positively correlated with *Sry* expression in the brain (Milsted et al., 2004; Dewing et al., 2006; Rosenfeld, 2017), we also investigated neurons expressing tyrosine hydroxylase (Th), the rate limiting enzyme for dopamine synthesis, in the AVPV. According to maternal behaviours and the link between sex chromosomes and vasopressin or Th expression in the brain, we expected a greater number of oxytocin, vasopressin and Th expressing neurons in the PVN and AVPV of X*Y females, respectively.

**Figure 1:**
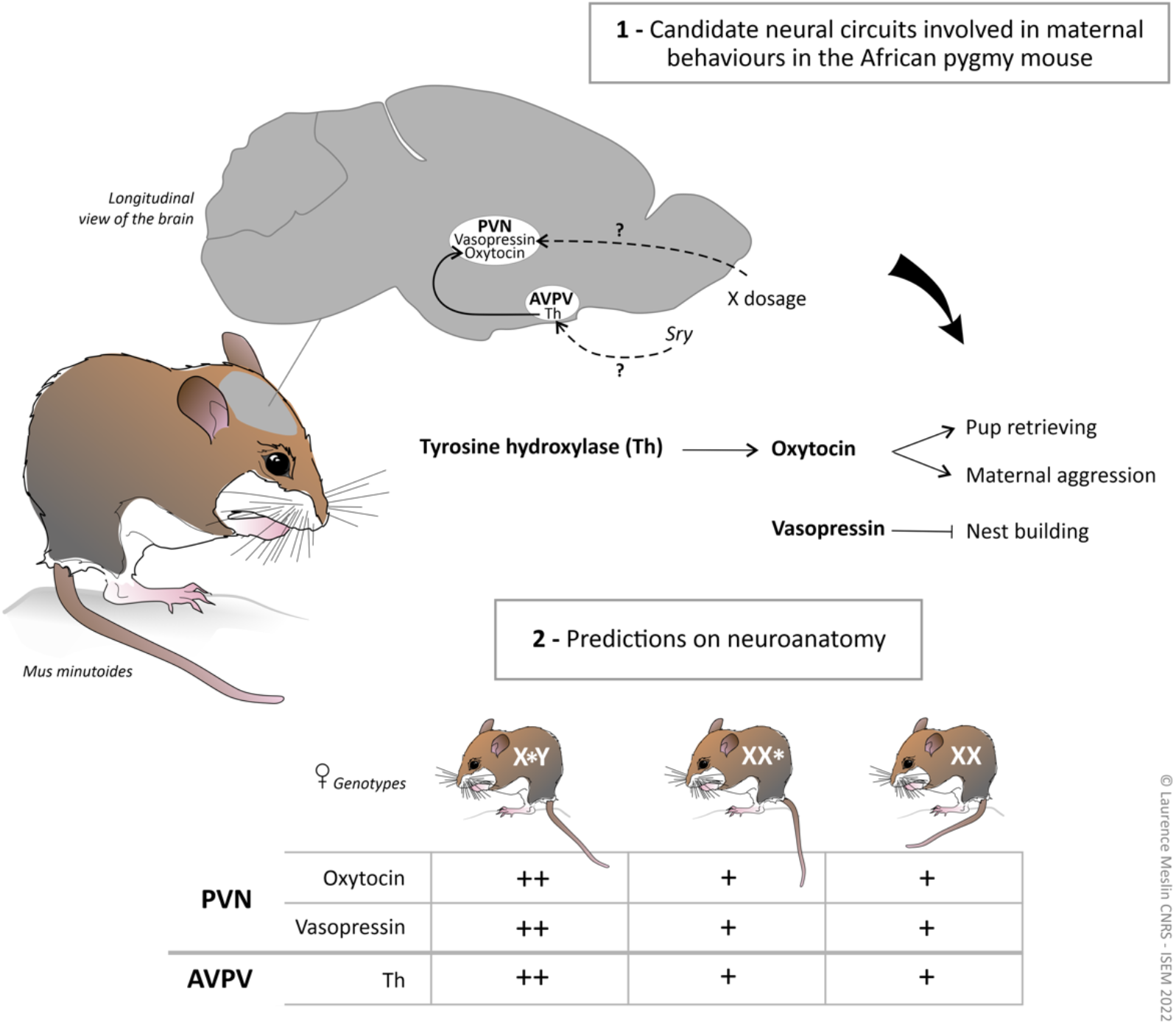
Neural investigations and predictions based on maternal behaviours in the African pygmy mouse as well as on the direct influence of sex chromosome on neuropeptides and neurotransmitters. Maternal behaviour experiments showed that sex-reversal in *M. minutoides* influenced pup retrieval, maternal aggression and nest building, with X*Y females being different from both XX and XX* females for each behaviour. We investigated potential candidates for the neural basis of these behaviours based on M. musculus studies: the paraventricular nucleus (PVN) of the hypothalamus where the expression of oxytocin triggers pup retrieval and maternal aggression (i.e. positive correlation) (Scott et al., 2015; Bosch & Neumann, 2012; Okabe et al., 2017; Pedersen et al., 2006) while vasopressin expression inhibits nesting behaviours (i.e., negative correlation) (Bendesky et al., 2017). We also investigated Th neurons (i.e., dopaminergic system) in the anteroventral periventricular nucleus (AVPV) of the hypothalamus, which have been shown to enhance retrieval behaviours through the regulation of oxytocin neurons in the PVN (Scott et al., 2015). Moreover, it has been shown that Th neurons can be regulated by the sex determining gene, Sry (e.g. Dewing et al., 2006) and that vasopressin can be influenced by X dosage (e.g. Gatewood et al., 2006). We thus predicted greater oxytocin, vasopressin and Th cell numbers / density in the PVN and AVPV of X*Y females, respectively. © Laurence Meslin CNRS -ISEM 2022

## Methods

### Animals, genotype identification and study environment

Behavioural experiments were conducted in accordance with European guidelines and with the approval of the French Ethical Committee on animal care and use (No. CEEA-LR-12170) in our laboratory colony (CECEMA facilities of Montpellier University) established from wild-caught animals from Caledon Nature Reserve (South Africa). Approximately 24 generations of mice were bred in our colony. Individuals are reared in large (38 L × 26 W × 24 H cm) or medium cages (31 L × 21 W × 21 H cm) filled with woodchips’ bedding, in a room held at 23 °C and with a 15:9 h light-dark cycle. They are supplied with food (mixed bird seeds) and water ad libitum. At weaning (i.e., six weeks post-parturition), virgin females are housed in groups of 5-6 in large cages. In our study, either females remained virgins or they were paired with a male at 2-10 months old in a medium cage for breeding and subsequent behavioural assays. Female genotype was identified whether by PCR of Sry using total genomic DNA extracted from tail tip biopsies with DNeasy blood and tissue kit (Qiagen) and following Veyrunes et al. (2010a) protocol or by karyotyping using bone marrow of yeast-stimulated individuals as previously described (Veyrunes et al., 2010b). We included X*Y and XX* females which were both littermated of either XX* or X*Y females, and XX females that were littermated of XX or XX* females.

### Behavioural assays

We tested primiparous females (i.e. first litter) aged 5.5 months old on average. Tests were conducted during the day, between 10.00 am and 6.30 pm, by an experimenter blind to females’ genotype. Each assay was filmed for subsequent analysis, with the exception of the parental care strategy test, which was scored by one-off observations in the cage. When individuals were assessed in more than one assay, we conducted experiments sequentially by increasing magnitude of stress: we first tested females for pup retrieval, followed by mother-pup interactions and then, nest building, with a 15 hours gap at least between each test. Behavioural data were analysed blindly regarding females’ genotype, using Observer v5.0.31 and Observer XT softwares (Noldus).

Pup retrieval experiments were carried out in home cage between 2-10 days post-parturition (dpp). The father and mother were removed while pup(s) (i.e., whole litter) were placed at the opposite corner of the shelter with the nest. An assay started once the female was put back under the shelter and recorded until the first pup was retrieved to account for litter size differences. If no pup had been retrieved within 10 minutes, the assay was ended. A total of 101 females (31 XX, 35 XX* & 35 X*Y) were assessed on both first approach latency and retrieval probability.

Mother-pup interactions experiments were carried out in a new cage (38 × 26 × 24 cm) filled with the individuals’ own bedding. We tested females between 2-10 dpp. We placed only one pup at the centre of the cage to account for litter size differences between females. We then placed the mother at a corner of the cage and recorded grooming, crouching and carrying behavioural responses during 10 minutes. We assessed 75 females (25 per genotype) on whether they displace/carry their pup, as well as on duration of grooming, huddling and crouching behaviours.

Nesting tests were performed in home cage, early evening and nest were scored in the following morning after 12 hours of experimentation for 90 females (34 X*Y, 27 XX and 29 XX*). Females were tested between 2-10 dpp. The father was removed from the cage as well as food, water, and all environmental enrichments except bedding. We added one nestlet (5×5 cm of compressed cellulose; Serlab, D00009) to provide nesting materials. Individuals received the same nestlet prior to experiment (post-parturition) for habituation. Nest scoring was assessed from 0 to 5 according to Gaskill et al. (2013) graduation: 0-cellulose unused and no nest, 1-no nest site, 2-flat nest, 3-slightly curved nest, 4-nest with walls, 5-fully enclosed nest. We scored the nest accordingly to the graduation above but with more flexibility allowing intermediary scores as well (e.g. 3.5 or 4.5). We also recorded the percentage of nestlet used by females as they do not necessarily use it to build the nest (i.e., they can use their bedding).

Parental care strategy, or whether mice display biparental care versus maternal care only, was investigated for 105 females (29 XX, 37 X*Y and 39 XX*) in home cage. We assessed strategies by characterizing fathers’ involvement with pup(s), the latter reflecting females’ behaviour towards them (i.e., fathers included or excluded from parental care). We made repeated one-off observations of their position in the cage and possible physical injuries, from birth to 14 dpp (before pup(s) start(s) to explore outside the shelter) and every 2-3 days. We rated fathers’ involvement gradually and increasingly, from 1 to 6: 1= father killed or removed for severe injuries, 2= chased, with aggression stigmata, 3= always in a different lodge, not tolerated in the shelter with the pup(s), 4= tolerated inside the shelter but in the opposite corner, 5= under the same shelter, next to the mother and 6= paternal care observed: huddling, carrying, crouching. Score measurements were not repeated if the father was killed or removed (i.e., score 1 or 2, referred to as high level of maternal aggression).

### Immunohistochemistry

We sampled virgin individuals only as no experimentations leading to death can be made on mothers. Eighteen virgin mice (3-5 per genotype including males as a control group) were euthanised with isoflurane 4% anaesthesia followed by decapitation. Brains were collected in 4% PFA and stored for at least 48 hours at 4°C. Brains were sliced at 30 μm using a vibratome (Leica Microsystems) and coronal sections were collected into a PBS-filled container. Floating brain slices were then put in 4×3 well plates (4 wells per individual) and washed three times with Phosphate Buffer Saline (PBS) before staining. We sliced and stained the whole brain to account for any potential divergence from M. musculus in the brain anatomy (no data were available on the brain of *M. minutoides*). Brain slices were first incubated for 24h at 4°C or two hours at room temperature (RT) with a Goat coupled Fab anti-mouse IgG blocking buffer (1:1000, Abliance) diluted in PBS with 0.1% Triton 100X (PBS.T) and ^~^1% BSA to prevent nonspecific binding. Then, slices were incubated for 24-48h at 4°C with mouse anti-Neurophysin I (1:2000, Merck Millipore, marker for Oxytocin), rabbit anti-vasopressin (1:1000, Immunostar) and sheep anti-Th (1:1000, abcam) diluted in PBS.T and BSA, followed by secondary antibodies : Cy3 coupled Donkey anti-mouse IgG antibody (1:1000, Jackson Laboratories), Cy5 coupled Donkey anti-rabbit IgG (1:1000,Jackson Laboratories) and A488 coupled Donkey anti-sheep IgG (1:1000,Jackson Laboratories) diluted in PBS.T and BSA, for 24-48 hours at 4°C. Slices were finally incubated for 3-6 minutes with DAPI (1:10 000, Invitrogen) diluted in PBS and mounted on Polysine® Slides stored at 4°C until imaging.

### Imaging and cell counts

Brain slices including the PVN and AVPV were identified based on the mouse brain atlas from Franklin and Paxinos (2001). Selected slices were processed with a scanner (Zeiss Axioscan) or confocal microscopy (Zeiss, LSM 780) and cell counts were done blindly and manually on imageJ, for entire nuclei. We also quantified neurons on a few brain slices using Imaris v9.8 software to compare with manual counts (only for oxytocin as the software did not perform well for vasopressin neurons). Oxytocin and vasopressin cell counts as well as their distribution across the whole PVN nucleus were investigated to account for potential differences in localization between genotypes.

### Statistical analysis

All statistical analyses were conducted on R v4.1.2 (R Core Team, 2021) using *lme4, stanarm, ordinal* and *multgee* packages (see Heitzmann et al., 2022a, 2022b for data and scripts, respectively). Pup retrieval was investigated with bayesian (generalized) linear mixed models, incorporating individuals’ age as a random effect on the intercept. We fit our models to assess retrieval probability and visit latency using a log(x+1)-transformation to correct for a right-skewed distribution (many latencies at 600s). We ran four Markov chains of 2000 iterations each, removed the first 1000 (warm-up) and kept the second half to estimate our parameters. Both our models checked convergence recommendations 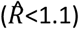. Post-hoc comparisons were made using *emmeans* package to contrast each genotype. Point estimates reported per genotype are medians (back transformed from the log (x+1) scale) and median-based contrasts with 89% Highest Posterior Density intervals (HPD) excluding one were considered as truly different. Mother-pup interactions were also analysed with Bayesian (generalized) linear mixed models with individuals’ age as a random effect and following the same steps as stated above. We log(x+1)-transformed duration responses to correct for a left-skewed distribution (many durations at 0s) and post-hoc comparisons were made to contrast genotypes. Nesting scores were analysed with a cumulative logit model (clm) and probabilities for nest quality per genotype were retrieved accordingly. XX was set as the reference group and we made multiple pairwise comparisons (*emmeans*) with adjusted p-value to assess each genotype effect. Parental care strategy was assessed with a population-averaged ordinal model to account for repeated measures and possible correlation between observations using the ordLORgee() function from *multgee* package (Touloumis, 2015). Probabilities were then retrieved using a back-transformation of parameters estimation according to a cumulative logit link. Cell counts for immunostaining assays were analysed using the non-parametric Kruskal Wallis test for multiple group comparisons.

## Results

### Higher retrieval efficiency in X*Y females

During pup retrieval experiments, X*Y females approached their pup(s) significantly faster than XX and XX* females (Fig. 2a, Fig. S1, table S1 in Heitzmann et al., 2022c), with a median visit latency at 51.7s [33.6-73.8 89% HPD] versus 156.5s [94.5-223.3 89% HPD] for XX and 112s [71.5-158.6 89% HPD] for XX* females. Interestingly, 35.5 % of XX females did not visit their pup(s) while only 17.1% of XX* females and 11.4% of X*Y did not respond (table S2). To confirm that differences were not driven by the absence of response, we reanalysed latencies only on females visiting their pup(s), and we found the same results (Fig S2). Additionally, X*Y females were significantly more likely to retrieve their pup(s) when compared to XX and XX* females, with an estimated retrieval probability of 0.87 [0.77, 0.95 89% HPD], while XX and XX* had a respective probability of 0.47 [0.32, 0.63 89% HPD] and 0.55 [0.42, 0.70 89% HPD] only (Fig. 2b, table S1 in Heitzmann et al., 2022c). Hence, X*Y females are more efficient at pup retrieval than XX and XX* females.

**Figure 2.**
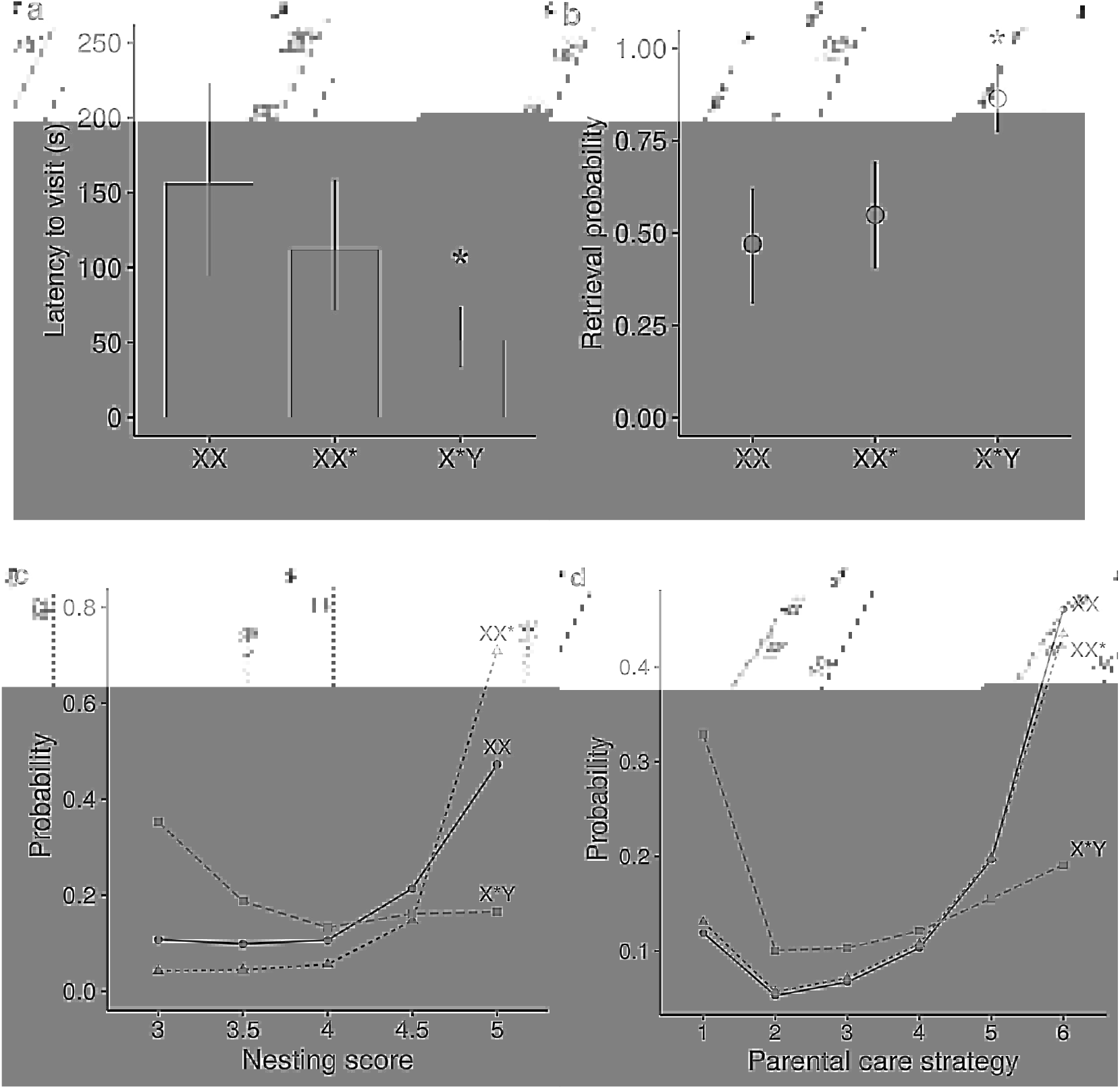
The sex chromosome complement shapes maternal behaviours in the African pygmy mouse. **a,b**, Pup retrieval. Estimates of first visit latency and retrieval probability per genotype. Values are medians with 89% HPD intervals. Latency to first visit and probabilities are retrieved from a back-transformation of our parameters. *Asterisks indicate significance of the genotype effect : X*Y females show a significantly reduced visit latency and a higher retrieval probability in contrast to XX as well as to XX* females (median-based contrasts with 89% HPD intervals are shown in table S2). nXX=31, nXX*=35, nX*Y=35 **c,** Likelihood curves of nest quality depending on females’ genotype. Nest quality estimates range from 3 (slightly curved nest) to 5 (fully enclosed nest) as no values below 3 were recorded. X*Y females build poorer quality nest than both XX (p=0.002) and XX* females (p<.0001). nXX=27, nXX*=29, nX*Y=34 **d,** Likelihood curves of parental care strategies regarding fathers involvement inside the nest from 1 : no involvement of fathers and high level of maternal aggression to 6 : biparental care. nXX=29, nXX*=39, nX*Y=37. X*Y females are more likely to display maternal aggression than both XX (p=0.005) and XX* females (p=0.004), which likely share care with males.

### Poor nest quality in X*Y females

All females showed nesting behaviours as no nest score below 3 was recorded. However, X*Y females were more likely to build poor nest quality in contrast to both XX (Fig. 2c, z= −3.09, p=0.006) and XX* females (z=−4.61, p< 0.001). They had a probability of 0.35 (highest probability) for building a slightly curved nest (i.e. score 3) while they had a 0.17 probability, for making a fully enclosed nest (i.e. score 5). Conversely, XX and especially XX* individuals were more likely to build high quality nests, with a respective probability of 0.47 and 0.71 for a fully enclosed nest. This also indicates a relatively high consistency in nest-building behaviour for XX* females. Though XX* had a higher probability than XX individuals for building high quality nests, they show similar behavioural trends (Fig. 2c, z= −1.82, p= 0.16). Interestingly, most X*Y females did not use the nestlet to build a nest (88.2 %) in contrast to 48.1 % for XX and 41.4% for XX*. We therefore tested for a relationship between nest quality and nestlet usage (i.e. disinterest for the nestlet or poor learning) in X*Y females and did not find any correlation (χ^2^ =10.05, p=0.19). X*Y females are thus less efficient at nest building when compared to XX and XX* females.

### Genotypic sex does not impact mother-pup interactions

Most females interacted with their pup and there were no significant differences in interaction durations nor probabilities (table S3, Fig. S3-4 in Heitzmann et al., 2022c). X*Y, XX and XX* females spent in median 7.9s [3.07-13.2 89% HPD], 11.2s [4.65-18.8 89% HPD] and 15.9s [7.03-25.8 89% HPD] crouching over the pup, 7.17s [3.48-11.36 89% HPD], 4.96s [2.26-7.97 89% HPD] and 7.30s [3.58-11.64 89% HPD] huddling with the pup, 3.18s [1.36-4.99 89% HPD], 4.87s [8.49-7.46 89% HPD] and 5.79s [2.89-8.70 89% HPD] grooming the pup, respectively. In addition, they had respective probabilities of 0.68 [0.42-0.9 89% HPD], 0.76 [0.5-0.93 89% HPD] and 0.67 [0.4-0.87 89% HPD] to displace their pup.

### Genotypic sex influences female parental strategy

X*Y females were on average more likely to show maternal aggression than XX and XX* females, which inversely, were more likely to be involved in biparental care (Fig. 2d, X*Y vs XX, z=-2.81, p=0.005; X*Y vs XX*, z=−2.87, p=0.004). Indeed, X*Y females had a highest probability at 0.33 to severely aggress or kill males (i.e., score 1) in contrast to a 0.19 probability to display biparental care (i.e., score 6). Inversely, XX and XX* females had highest probabilities at 0.46 and 0.43 respectively for biparental care in comparison to 0.12 and 0.13 probabilities for maternal aggression. Strikingly, XX and XX* females’ curves almost overlay with quite similar probabilities in each category highlighting identical parental care strategy in XX and XX* females.

### Hypothalamic neural circuits of maternal care

We did not find evidence for difference in the global neuroanatomy of *M. minutoides* PVN in comparison to *M. musculus* (Gimpl & Fahrenholz, 2001; Franklin & Paxinos, 2001; Madrigal & Jurado, 2021).Vasopressin is expressed in both parvo- and magnocellular neurons, mostly in intermediate regions of the PVN where it is expressed dorsolaterally (Fig. S5, S7-S10 in Heitzmann et al., 2022c) and oxytocin is mostly expressed in magnocellular neurons and more homogeneously across the nucleus than vasopressin (midline and rostro-caudal axis; Fig. S6-S10 in Heitzmann et al., 2022c). We did not have enough power to statistically compare oxytocin and vasopressin neuron distributions across genotypes but there were no overall striking differences (Fig. S5-S6 in Heitzmann et al., 2022c), with high variabilities in count data due to inter-individual variability and small sample sizes. In addition, there were no differences in oxytocin neurons number between genotypes (Fig. 3a,c-S6, χ2 = 0.89, p= 0.83). We did not find differences in the number of vasopressin expressing neurons either (Fig. 3b,c, χ2=0.82, p= 0.84) but there is a tendency for lower number in X*Y females driven mostly by the lower number of vasopressin neurons in anterior regions (Fig. S5 in Heitzmann et al., 2022c). Finally, there were no differences in the neuroanatomy of the AVPV in comparison to *M. musculus* (Franklin & Paxinos, 2001; Simerly et al., 1985; Semaan & Kauffman, 2010). Since there is evidence for a sexual dimorphism in the nucleus size in *M. musculus* (Bleier et al., 1982; Semaan & Kauffman, 2010), we weighted the number of Th expressing neurons by the area of the nucleus and did not find differences in the density of Th neurons between genotypes (χ2=2, p= 0.57), though a tendency for a greater density in X*Y females (Fig. 4).

**Figure 3.**
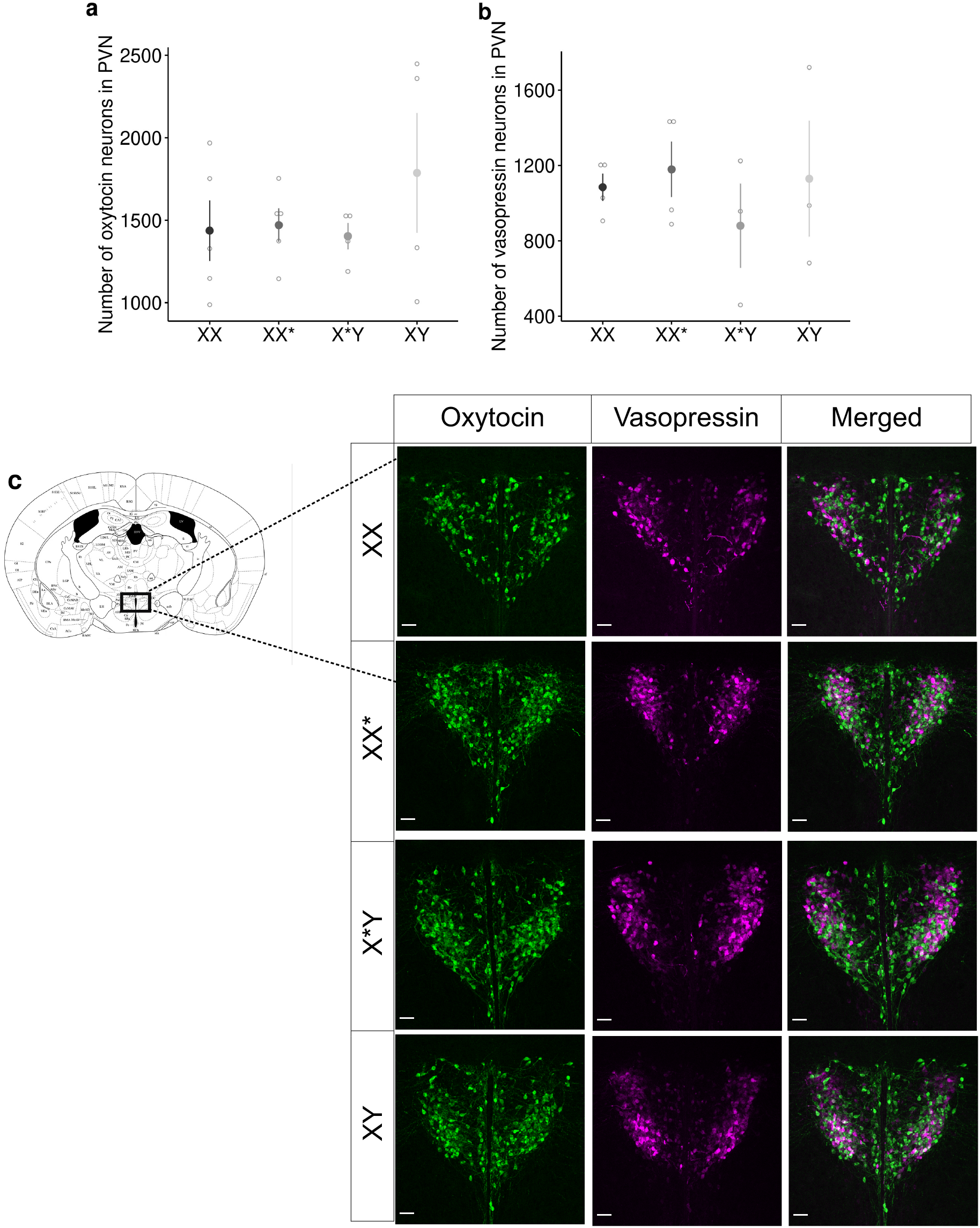
No genotypic sex effect on oxytocin and vasopressin neurons’ number in the PVN of virgin individuals. a,b, Total number of oxytocin and vasopressin expressing neurons in the PVN of virgin XX, XX*, X*Y females and XY males. Oxytocin : nXX=5, nXX*=5, nX*Y=4, nXY=4. Vasopressin : nXX=4, nXX*=4, nX*Y=3, nXY=3. Data are means ± s.e.m and raw counts are also represented by empty dots. c, Oxytocin and vasopressin staining in the PVN of virgin African pygmy mice and associated M.musculus brain slice reference (black rectangle represents the PVN area, modified from Franklin & Paxinos, 2001). Scale bars 50 μm.

**Figure 4.**
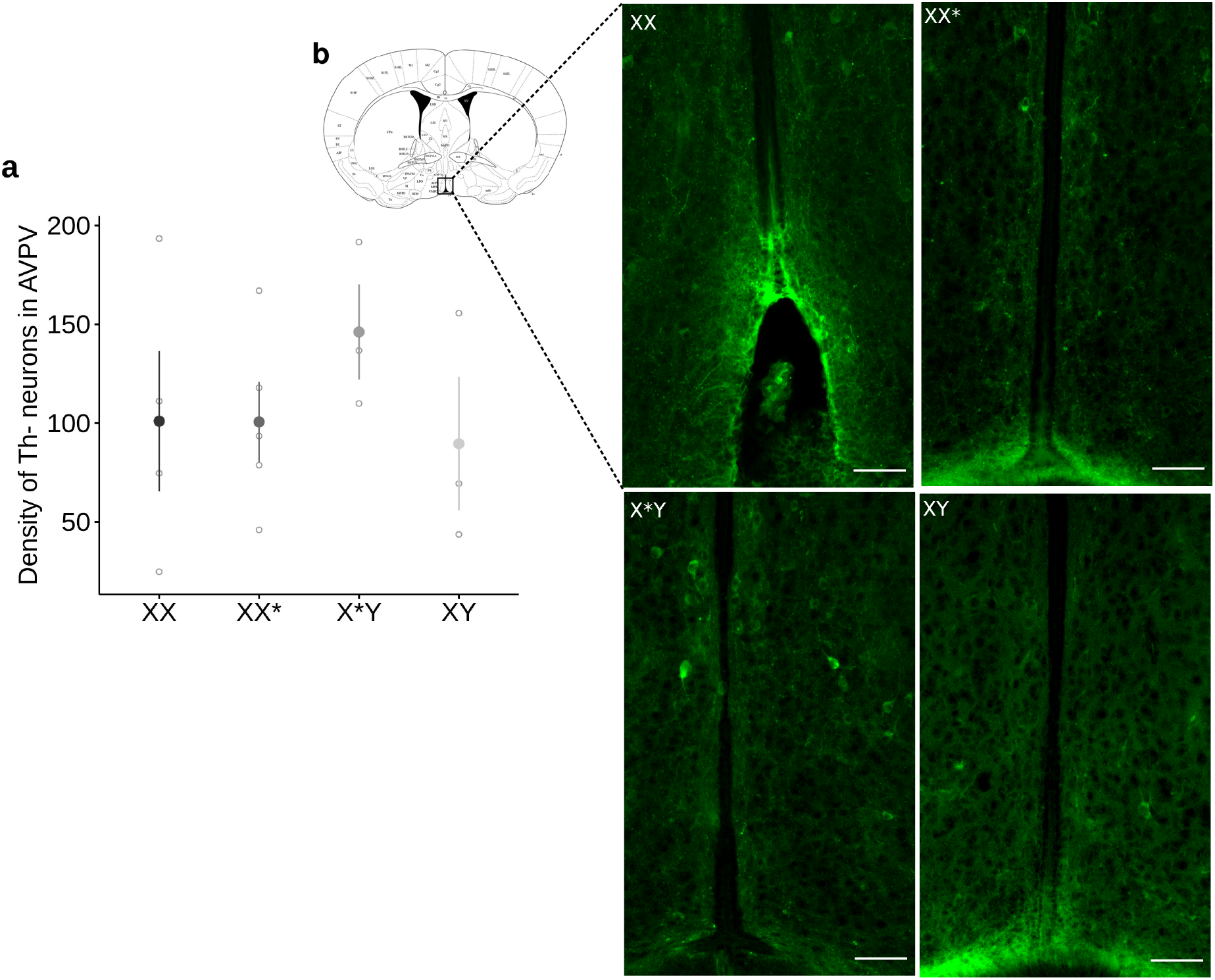
Tendency for greater Th neuron density in the AVPV of X*Y females. a, Density of Th-expressing neurons in PVN of virgin XX, XX*, X*Y females and XY males: number of total Th neurons weighted by the sum of the area (mm2). nXX=4, nXX*=5, nX*Y=3, nXY=3. Data are means ± s.e.m and raw counts are also represented by empty dots. b, Th staining in the AVPV of virgin African pygmy mice and associated *M.musculus* brain slice reference (black rectangle represents the AVPV area, modified from Franklin & Paxinos, 2001). Scale bars 50 μm.

## Discussion

The African pygmy mouse is a species with a naturally occurring sex reversal system, involving a dissociation between phenotypic and genotypic sexes in X*Y females. This fascinating system represents a great opportunity to shed light on the impact of the sex chromosomes on sexually dimorphic behaviours such as parental care. By focusing on maternal phenotypes specifically, we show that sex reversal in *M. minutoides* influences pup retrieval, maternal aggression, nest building behaviours and may also impact the maternal dopaminergic circuit in the AVPV nucleus. Our study confirms and further extends evidence on the influence of the sex chromosome complement on parental behaviours and brings promising prospects for its impact on the neurobiology of parenting.

X*Y females are more efficient at pup retrieval than both XX and XX* females. They first approached their pup faster and showed more consistent retrieval, while XX and XX* showed delayed latencies to first approach and retrieve their pup about one out of two times. Moreover, in a reproductive context, X*Y mothers were highly aggressive towards males, whereas XX and XX* females were more likely to share parental care. Previous results showed that non-reproductive X*Y females are more aggressive than other female genotypes towards intruder males (Saunders et al., 2016). Here, we show this enhanced aggressiveness further extends towards the pup’s own father (as male partners were killed or needed to be removed for 43.2% of X*Y females (i.e., 16 out of 37)), suggesting that X*Y females are more likely to be solitary mothers. Furthermore, X*Y females built poor quality nests (slightly curved), while XX and especially XX* females performed better with fully enclosed nests. It was surprising that almost all X*Y females did not use the nestlet, and the lack of correlation between nestlet usage and nest quality strongly suggests a disinterest for the nestlet in X*Y females. Because nest building is an essential component of maternal care especially in altricial species, to protect the young from potential threats for instance, this would thus suggest maladaptation in X*Y females. However, we show here that X*Y females can assure pups protection through agonistic behaviours (i.e. maternal aggression), which might thus make the nest quality less essential for these females. Hence, the divergent behavioural patterns between X*Y and XX / XX* females reflect two distinct maternal strategies shaped by the sex chromosome complement in *M. minutoides*. Although there is an unequivocal effect of the sex chromosome complement on some maternal behaviours, there were no differences in mother-pup interactions suggesting that it does not modulate all component of maternal care.

It is very striking that X*Y females displayed -quite consistently-, divergent maternal behaviours from XX and XX* females: they showed a greater retrieval efficiency, high level of maternal aggression but also poor nesting skills as found in males suggesting masculinization (data not shown; Topilko et al., 2022). Interestingly, this pattern of co-occurrence of both feminized and masculinized traits in *M. minutoides* X*Y females was already found in previous studies. While X*Y females have functional ovaries and show moreover, a greater reproductive success than both XX and XX* females (e.g. higher ovulation rate, bigger litter sizes) (Rahmoun et al., 2014; Saunders et al., 2014), they also display male-like reduced anxiety behaviours, they are more aggressive, and have a greater bite strength than XX and XX* females (Saunders et al., 2016; Ginot et al., 2017). All things considered, the constant divergence between X*Y on one side and XX and XX* females on the other side highlights that the association of a feminizing X* and a male-specific Y chromosome has given birth to a third sexual phenotype in the African pygmy mouse.

The differences of maternal care behaviours between female genotypes raise questions concerning the genetic bases of maternal strategies in *M. minutoides*. Several hypotheses are discussed here. First, the divergent behaviours of the X*Y females suggest an influence of the Y chromosome and one prime candidate among this gene-poor chromosome is *Sry*, which is expressed in the brain of X*Y females (Rahmoun et al., 2014). Secondly, there could also be an influence of the female specific X* chromosome. However, since XX* females behave like XX females, this would imply maternal X*-biased chromosome inactivation (X*CI), or parental imprinting of X-linked genes (Gregg et al., 2010; Raefski & O’Neil, 2005), with preferential expression of the paternal × alleles.

There is no evidence for skewed X*CI in *M. minutoides* based on cytological investigations of embryos fibroblast cell cultures (Veyrunes & Perez, 2018), but next generation sequencing technologies with a special focus on allele-specific expression in the brain would be necessary to confirm or refute this finding, and therefore help clarify hypotheses. Thirdly, another hypothesis to correlate with divergent maternal strategies is X dosage (i.e., copy number of X chromosomes). In mice, a thousands of autosomal genes are under the influence of X dosage (Wijchers et al., 2010) such as vasopressin, involved in nesting behaviours (Bendesky et al., 2017; De Vries et al., 2002; Gatewood et al., 2006; Cox et al., 2015). In addition, a few X-linked genes, such as Kdm6a involved in gene expression regulation, escape inactivation in mice and have also been suggested to influence sexual phenotypes (3-8%; Tsuchiya & Willard, 2000; Ngun et al., 2014; Arnold, 2020). Finally, on might also suppose an epistasis interaction between the X* and the Y chromosome. In Drosophila melanogaster for instance, it has been shown that a few X-linked genes involved in pheromone detection are regulated by the Y chromosome, which is likely to impact reproductive phenotypes (Jiang et al., 2010). Interestingly, some of our behavioural observations are in contradiction with previous findings on the FCG, which may give insight into the genetic bases. Contrary to X*Y females in *M. minutoides*, sex-reversed XY-females in the FCG displayed similar nesting behaviours to XX females but impaired retrieval capabilities: they retrieved less pups and belatedly when compared to XX females (Gatewood et al., 2006). Considering nest building, because the authors controlled for gonadal hormones, this suggested that the Sry gene and/or male gonadal hormones acting perinatally to masculinize brain neural circuits, were responsible for the poor nesting skills in males (Gatewood et al., 2006). In our case, X*Y females built poor quality nests while they do not have differences in gonadal/adrenal hormonal levels compared to the other female genotypes (testosterone and estradiol; Veyrunes et al., 2022) but they do express Sry in their brain (Rahmoun et al., 2014). Therefore, Sry may also influence nesting behaviours in the African pygmy mouse. Considering pup retrieval, results in the FCG suggested that the Y chromosome and/or X dosage imbalance was necessary to alter retrieval phenotypes. Here, we show that X*Y females perform better at pup retrieval even though they have one copy of the X* chromosome, a Y chromosome, and express Sry in their brain (Rahmoun et al., 2014). Therefore, either Y-linked genes including Sry act distinctly on neural pathways between FCG mice and this wild mouse species, either it is an X* chromosome effect or an epistasis interaction between the X* and the Y chromosome. Although the genetic bases with the relative part of X, X* and Y chromosomes in driving these divergent maternal behaviours requires further investigations, our results show that sex reversal in the African pygmy mouse does not seem to alter maternal care quality, contrary to expectations based on the FCG. This discrepancy further illustrates the need to consider not only genetically modified laboratory mice but also non-model species with alternative phenotypes shaped by natural selection -such as observed in *M. minutoides-*, to better understand the genetic and neural bases of behaviours.

In mammals, oxytocin and vasopressin are crucial neuropeptides that intervene in the initiation and maintenance of parental care (Zilkha et al., 2017). In rodents, oxytocin expression in the PVN induces pup retrieval and maternal aggression (Neumann et al., 2005 ; Pedersen et al., 2006; Scott et al., 2015; Okabe et al., 2017; Bosch & Neumann, 2012) while vasopressin, which can also mediate maternal aggression, has been shown to inhibit nest building (Bendesky et al., 2017; Bosch & Neumann, 2012). Moreover, vasopressin has been shown to be influenced by X chromosome number, with greater expression in individuals with a single X chromosome (De Vries et al., 2002; Gatewood et al., 2006, Cox et al., 2015). Hence, oxytocin and vasopressin were prime candidates to explain the divergent maternal behaviours in *M. minutoides*, and a greater number of vasopressin and oxytocin neurons in the PVN of X*Y females in comparison to XX and XX* females could have been expected (Fig. 1). However, we did not find such differences between females nor with males, emphasized by high variability in count data (i.e. indicative of low power; Fig. 3a,b). The lack of differences in oxytocinergic and vasopressinergic systems global anatomy in the PVN suggests no differences in oxytocin and vasopressin levels. Nonetheless, larger sample sizes and the functionality of the neuroendocrine networks as well as the hormonal output should be tested to confirm or refute this hypothesis (e.g. quantification of gene and protein expression in the PVN, blood hormonal levels measured over time) (Cox et al., 2015; Scott et al., 2015; Bendesky et al., 2017). In addition, we tested virgin females while parental neural circuits are known to evolve with sexual experience (Lopatina et al., 2011; Scott et al., 2015; Okabe et al., 2017; Zilkha, Scott & Kimchi, 2017). It is thus likely that differences in oxytocin and/or vasopressin between females arise at first pregnancy.

Another interesting brain region to look at was the AVPV. It has been shown that neurons expressing Tyrosine hydroxylase (Th) in the AVPV (AVPV^Th^) stimulate oxytocin expressing neurons in the PVN and consequently pup retrieval in females (Scott et al., 2015). Moreover, these neurons are sexually dimorphic with a greater number in females than males already in virgin individuals (Semaan & Kauffman, 2010; Scott et al., 2015). Our results do not support the sexual dimorphism described in *M. musculus*, however we show a tendency for a greater density of Th neurons in X*Y females, which is also supported by preliminary results on Th mRNA expression levels (RNAseq and RT-qPCR) in the whole brain of *M. minutoides* (unpublished data), as well as with the link between *Sry* and Th (Milsted et al., 2004; Dewing et al., 2006; Rosenfeld, 2017). Therefore, the tendency for Th may also be consistent with pup retrieval results. Though not significant, it brings new perspectives on the link between sex chromosomes and the dopaminergic system as well as between dopamine and parenting.

Finally, it is noteworthy that parenting is a complex behaviour that is regulated by diverse neural pathways and neuromodulators. In rats for instance, it has been shown that overexpression of the neuropeptide Y, which is sexually dimorphic, correlates with impaired nesting behaviours (Nahvi & Sabban, 2020 ; Corder et al., 2020). It is thus not unlikely that the sex chromosome complements in *M. minutoides* act on various neural circuits to shape maternal behaviours, and future studies should integrate other brain regions and neuromodulators, to help understand the neural basis of maternal strategies. Because we work on a wild non-model species, we are facing interpretation issues due to small sample sizes, the use of virgin individuals, and technical limitations (stereotaxic surgery has never been performed on this species), which restrain the establishment of a causal link between brain nucleus, peptides and behaviours. Nonetheless, primary investigations on the neuroanatomy are essential to describe and further understand the evolution of the brain and behaviours as illustrated here in a wild species with highly contrasted phenotypes.

Combining behavioural ecology and the neuroanatomy of parenting, we highlight the emergence of a third sexual phenotype in a wild species. While the relative impact of each sex chromosome on maternal behaviours is yet to be determined, our results bring exciting advances to research on both the neural basis and the impact of the sex chromosomes on natural behaviours. Furthermore, research on multiple sexual phenotypes in other wild species with unusual sex determination system as described here, in the African pygmy mouse, could help strengthen the challenge of rethinking the stereotypical dichotomous view of sexes.

## Acknowledgements

We thank Dr Paul Saunders for insightful comments, advices, and review on the manuscript; Dr Stanislav Cherepanov for oxytocin antibody and insight on the study; Anne Guillou for interesting discussions on the study and the model. We thank Caroline Hu, Marion Anne-Lise Picard and an anonymous reviewer for fruitful reviews and comments on the manuscript.We thank the animal breeding facility of the University of Montpellier (CECEMA), the imaging facility MRI and Laurence Meslin for the illustration. Preprint version 4 of this article has been peer-reviewed and recommended by Peer Community In Evolutionary Biology (https://doi.org/10.24072/pci.evolbiol.100152)

## Data, scripts, code, and supplementary information availability

Data are available online: https://doi.org/10.5061/dryad.c2fqz61b5

Scripts and code are available online: https://doi.org/10.5281/zenodo.7234022

Supplementary information is available online: https://doi.org/10.5281/zenodo.7156308

