## Supplementary Material for "Genotypic sex shapes maternal care in the African Pygmy mouse, *Mus minutoides*"

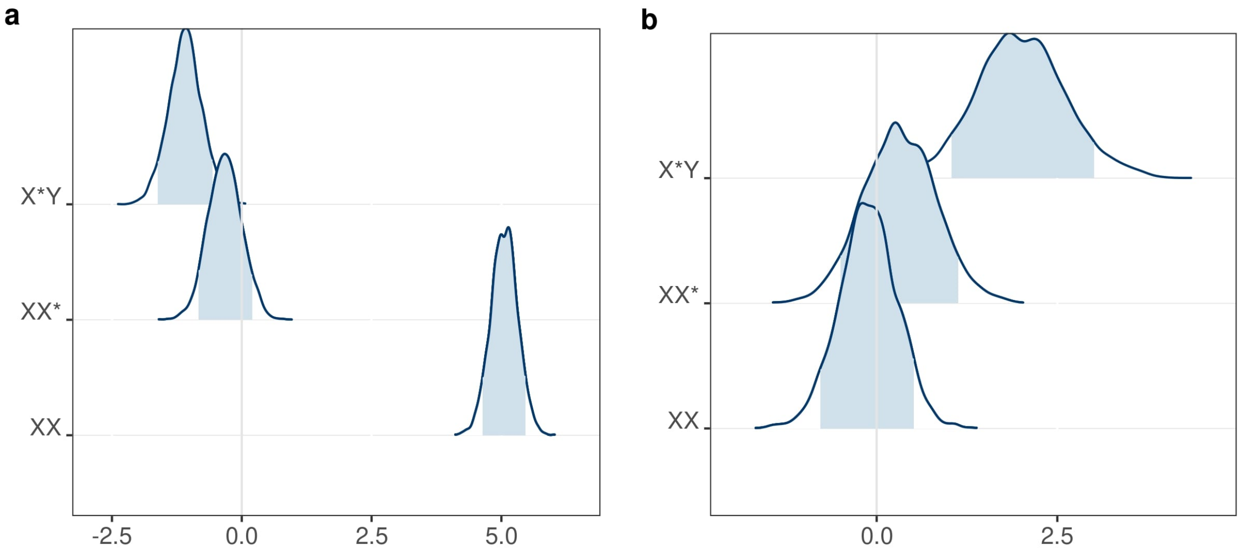


**Figure S1.** Posterior density interval estimates of genotype effect on pup retrieval. **a,b,** latency to first visit and retrieval probability respectively. XX females are defined as the reference population and slope estimates are represented for XX* and X*Y females effects. Estimates are on the log (x+1) and logit scale (**a** and **b** respectively). Shaded areas represent 89% Highest Posterior Density intervals (HPD) and groups whose HPD intervals cross the vertical grey line are considered as not significantly different from the reference group (XX).

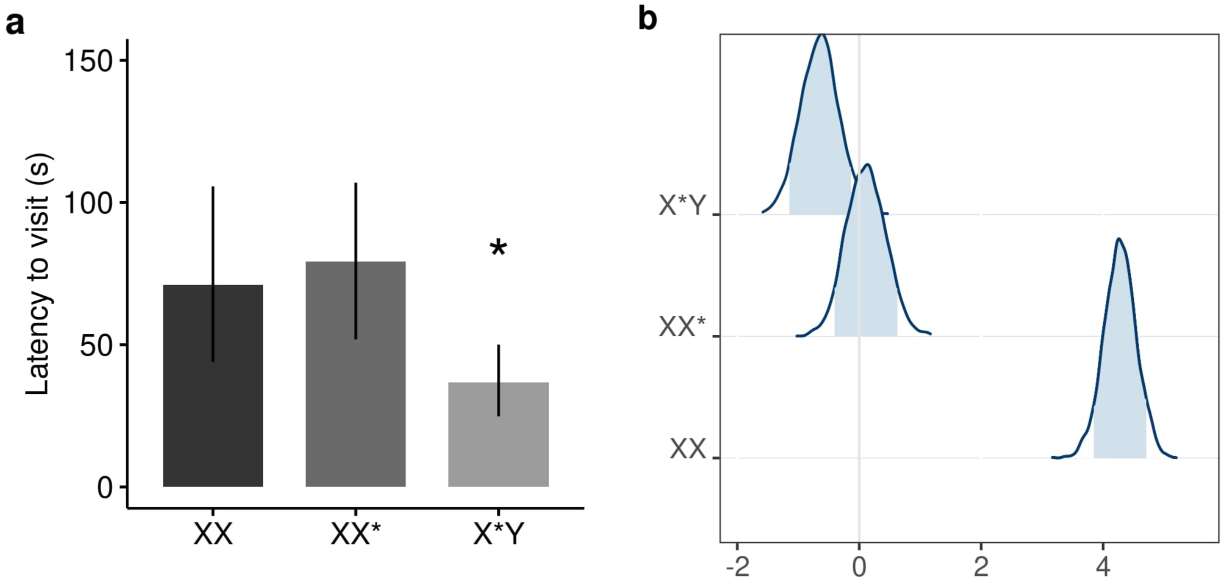


**Figure S2.** Visit latencies for females responding only. **a,** Estimates of first visit latency per genotype. Values are medians ± 89% HPD intervals. Estimates are retrieved from a back-transformation of models' parameters. *Asterisk indicate significance of the genotype effect. **b**, Posterior density interval estimates of genotype effect. XX females are defined as the reference population and slope estimates are represented for XX* and X*Y female effects. Estimates are on the log (x+1) scale. Shaded areas represent 89% Highest Posterior Density intervals (HPD) and group whose HPD intervals cross the vertical grey line are considered as not significantly different from the reference group (XX).$n_{XX}$=20 , $n_{XX*}$=29, $n_{X*Y}$=31.

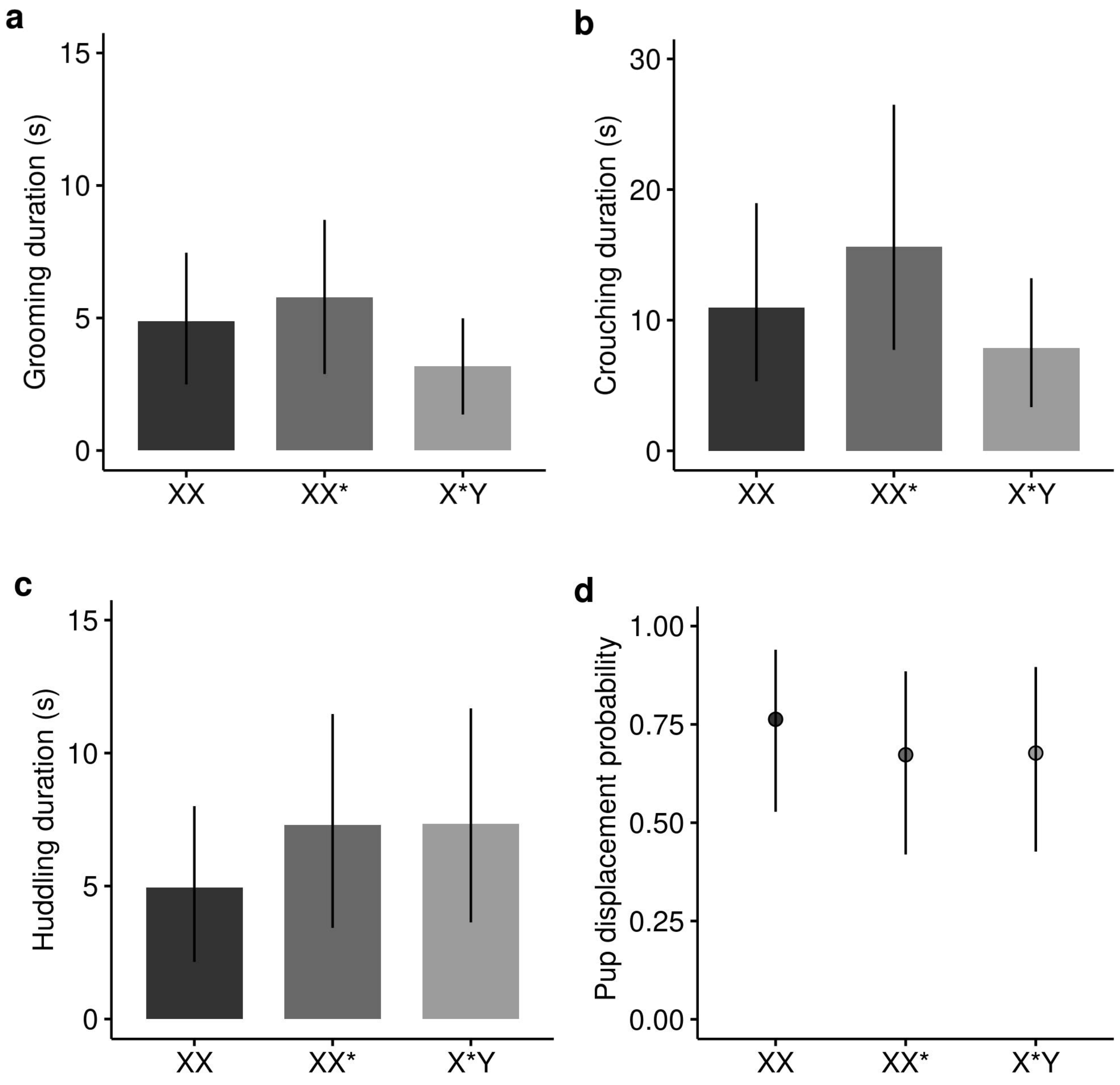


**Figure S3.** The sex chromosome complement does not influence direct mother-pup interactions in the African pygmy mouse. Values are medians ± 89% HPD intervals. Estimates are retrieved from a back-transformation of models' parameters. n=25 per genotype.

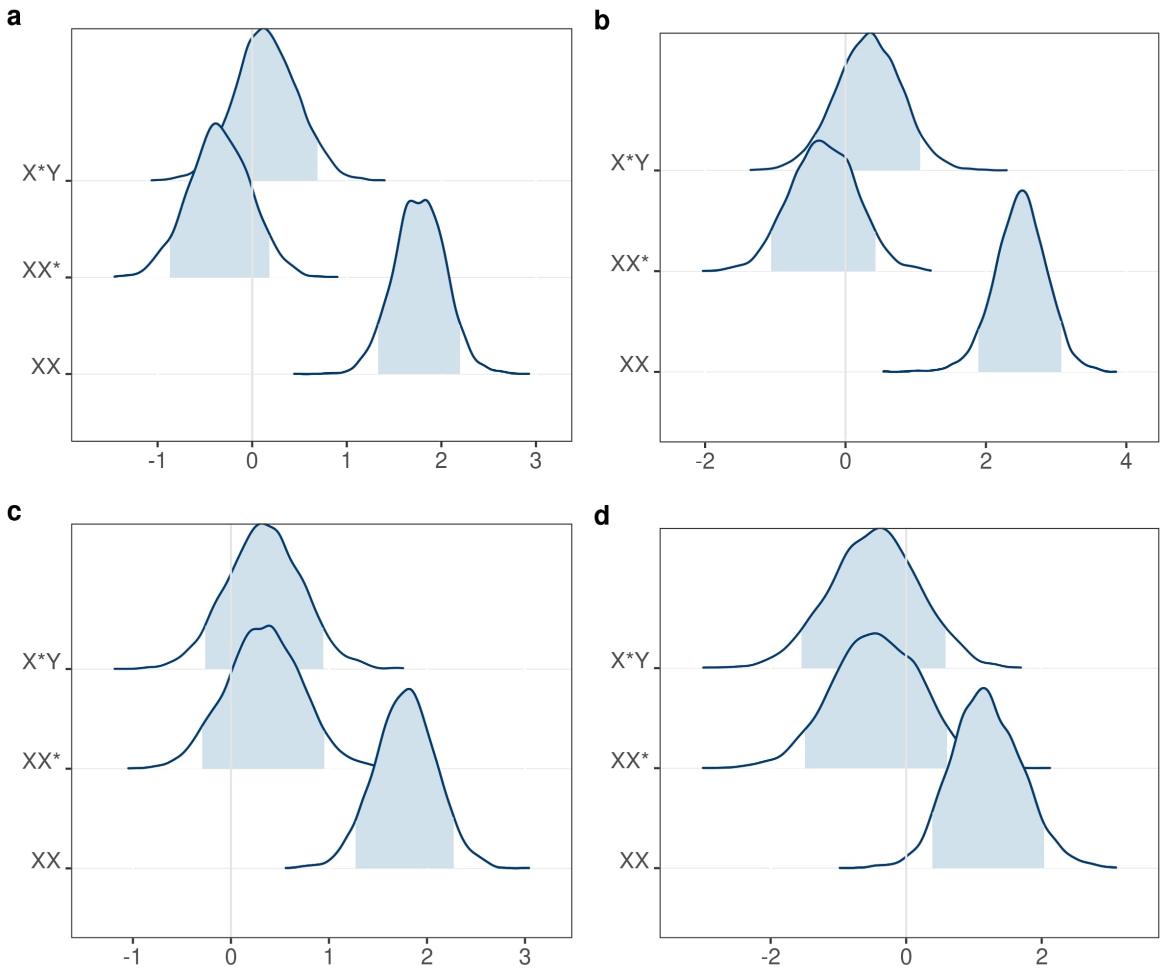


**Figure S4.** Posterior density interval estimates of genotype effect on mother-pup interactions. **a,** grooming duration. **b,** crouching duration. **c,** huddling duration. **d,** pup displacement in the cage probability. XX females are defined as the reference population and slope estimates are represented for XX* and X*Y females effects. Estimates are on the log (x+1) and logit scale (**a,b,c** and **d** respectively). Shaded areas represent 89% Highest Posterior Density intervals (HPD) and group whose HPD intervals cross the vertical grey line are considered as not significantly different from the reference group (XX). n=25 per genotype.

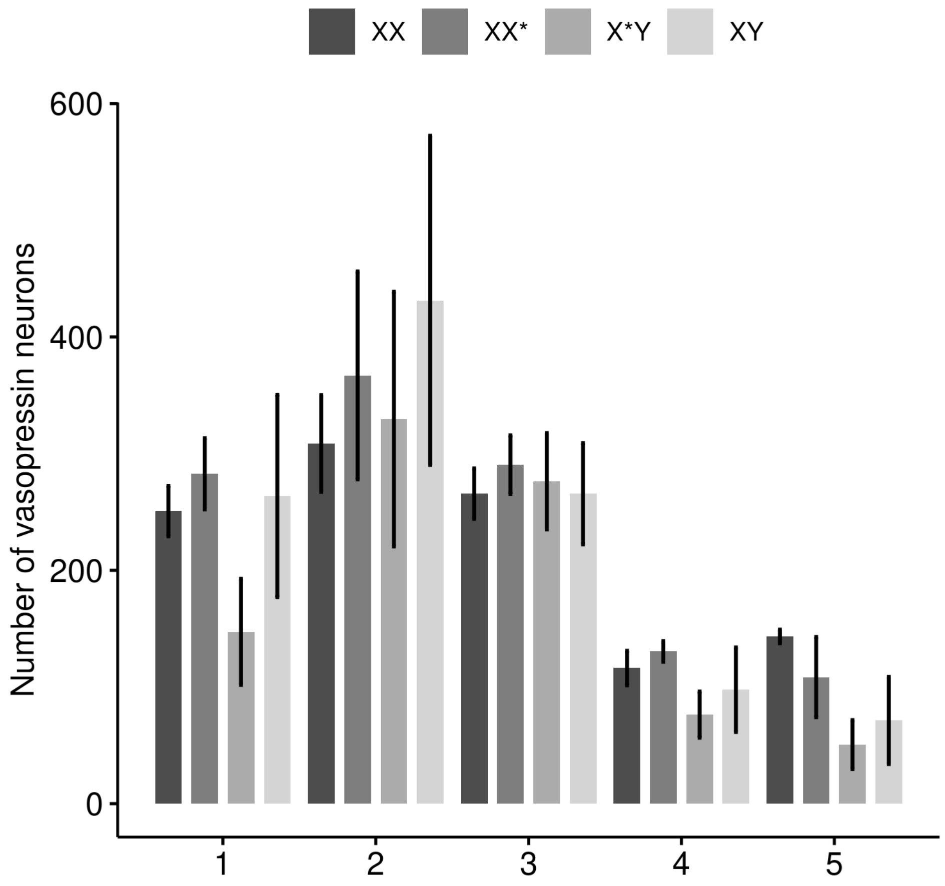


**Figure S5**. Vasopressin neurons distribution across the PVN. Values are mean ± s.e.m. We removed one X*Y female due to staining issues on few brain slices. Regions (1-5) are illustrated in Fig S7-S10. $n_{XX}$=4 , $n_{XX*}$=4, $n_{X*Y}$=2, $n_{XY}$=3. Males tend to have greater vasopressin neurons in region 2 (early intermediate region) and X*Y females tend to have lower vasopressin neurons in region 1 (anterior region).

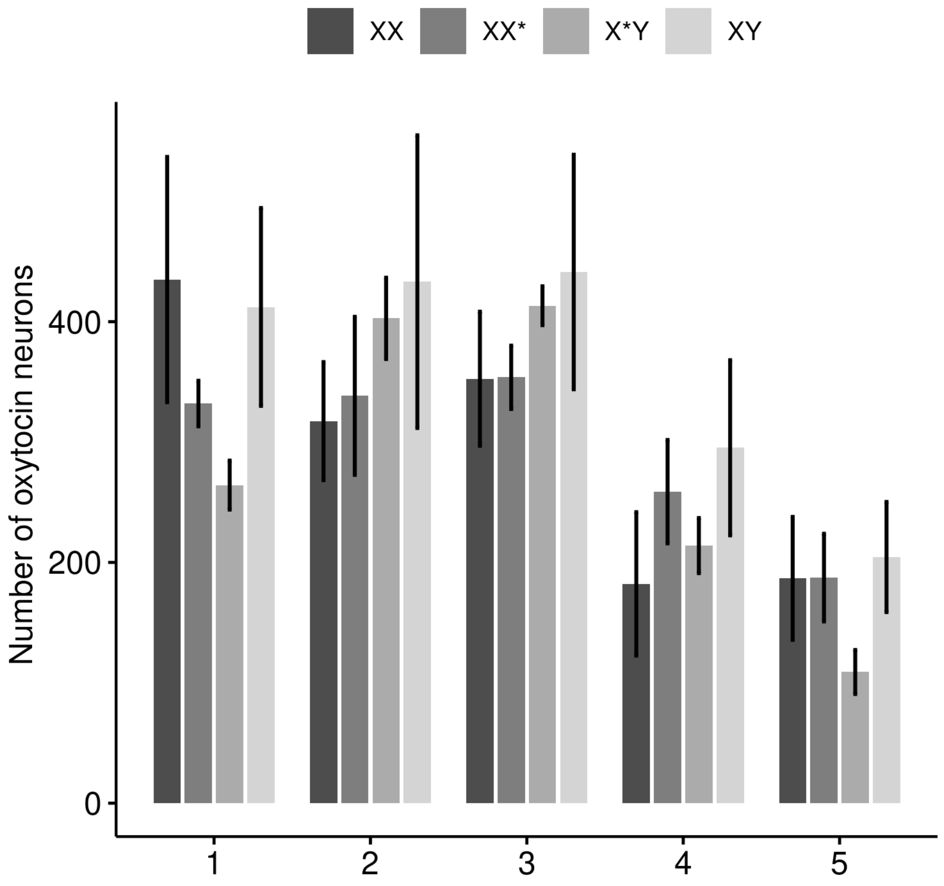


**Figure S6**. Oxytocin neurons distribution across the PVN. Values are mean ± s.e.m. Regions (1-5) are illustrated in Fig S7-S10. $n_{XX}$=5 , $n_{XX^*}$=5, $n_{X^*Y}$=4, $n_{XY}$=4.There is a tendency for a left skewed oxytocin distribution in XX females with a greater neuron number in anterior regions, while in intermediate regions for X*Y females and both regions for XX* and XY individuals. X*Y females tend to have a lower number of oxytocin neurons in region 1 and 5 in comparison to other individuals.

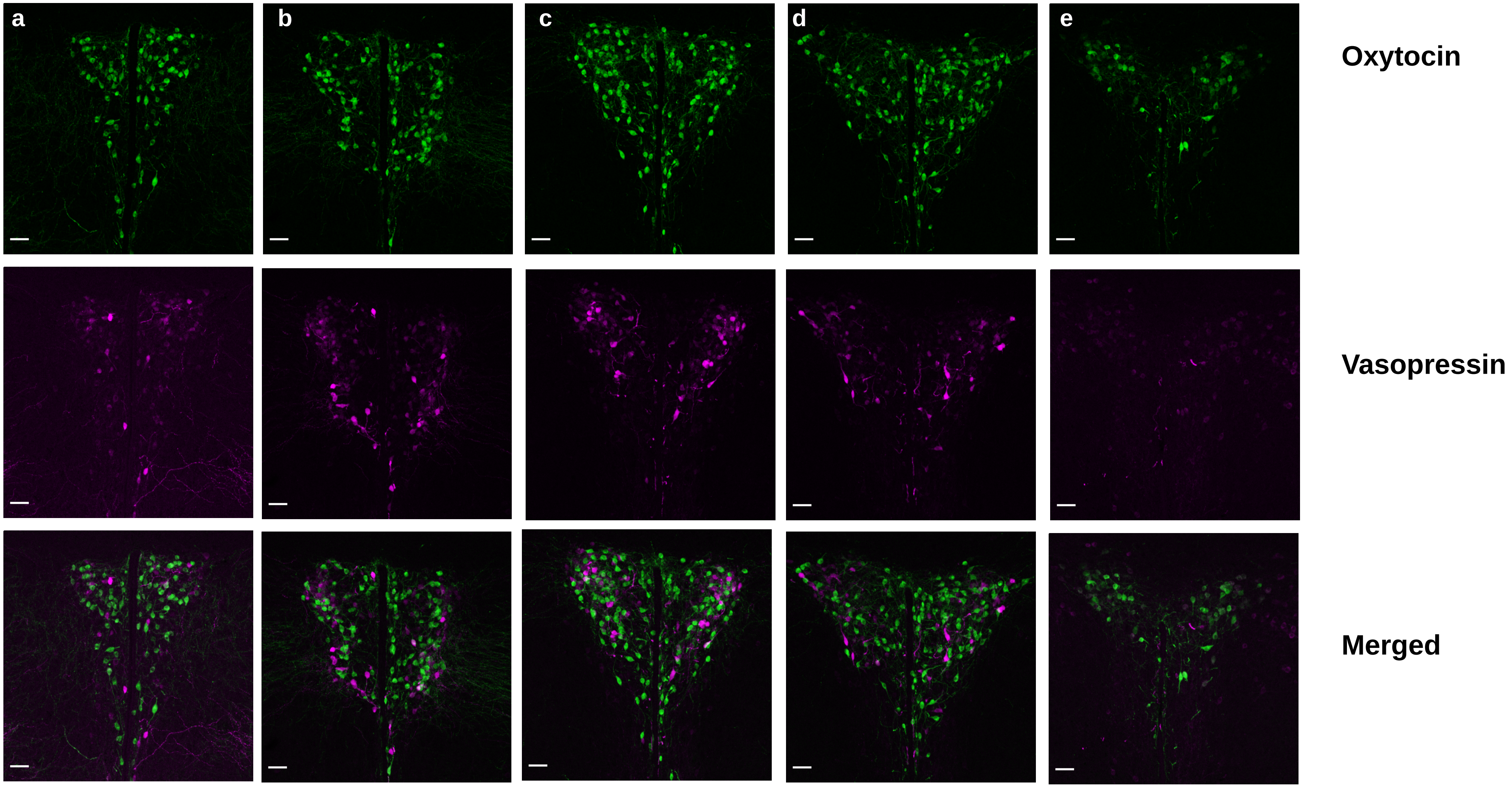


**Figure S7.** Oxytocin and vasopressin neurons' expression across the PVN in XX females. **a,** region 1 : anterior. **b,c**, regions 2 and 3 : intermediate. **d,e,** regions 4 and 5 : posterior.

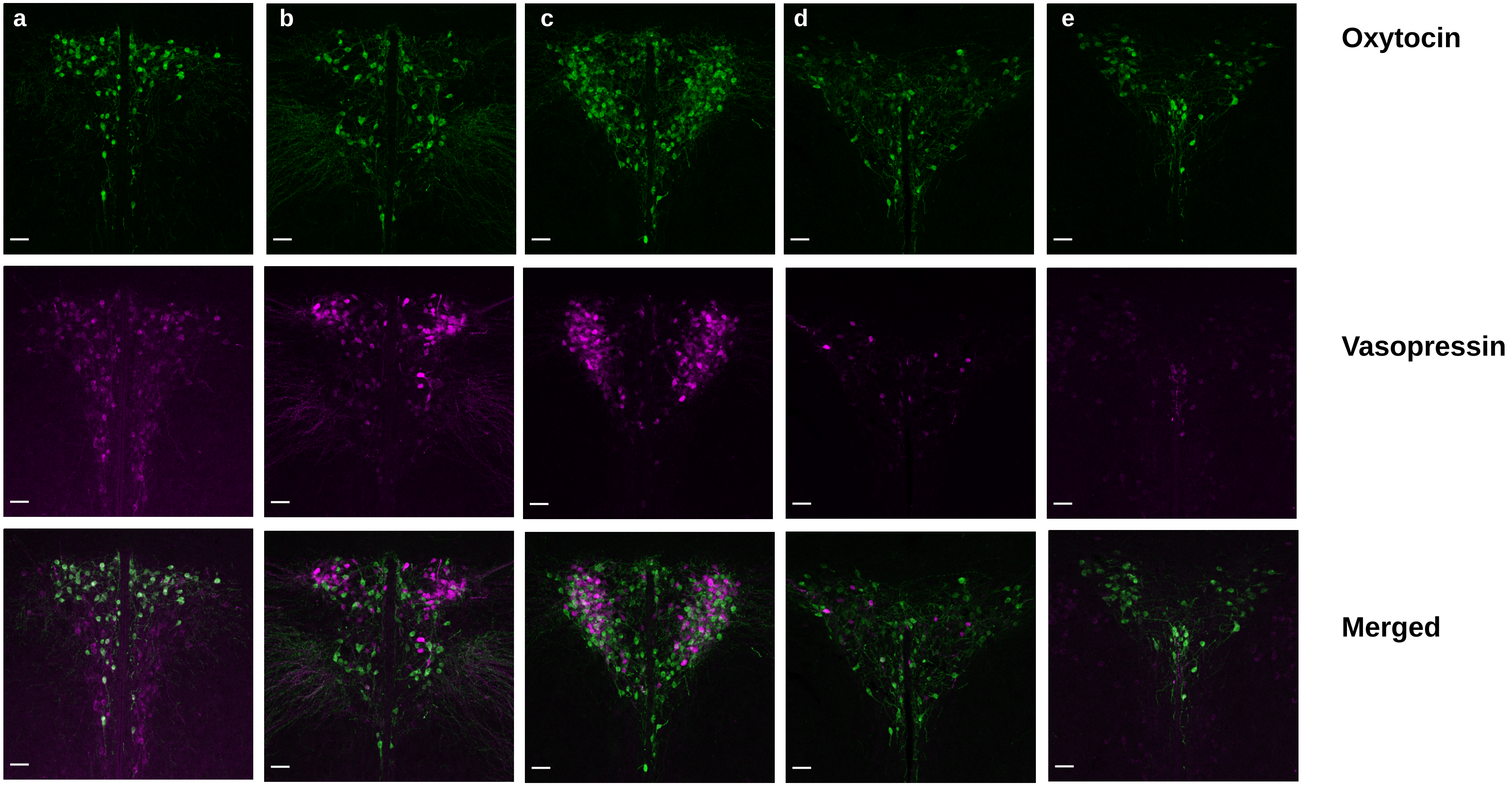


**Figure S8.** Oxytocin and vasopressin neurons' expression across the PVN in XX* females. **a,** region 1 : anterior. **b,c**, regions 2 and 3 : intermediate. **d,e,** regions 4 and 5 : posterior.

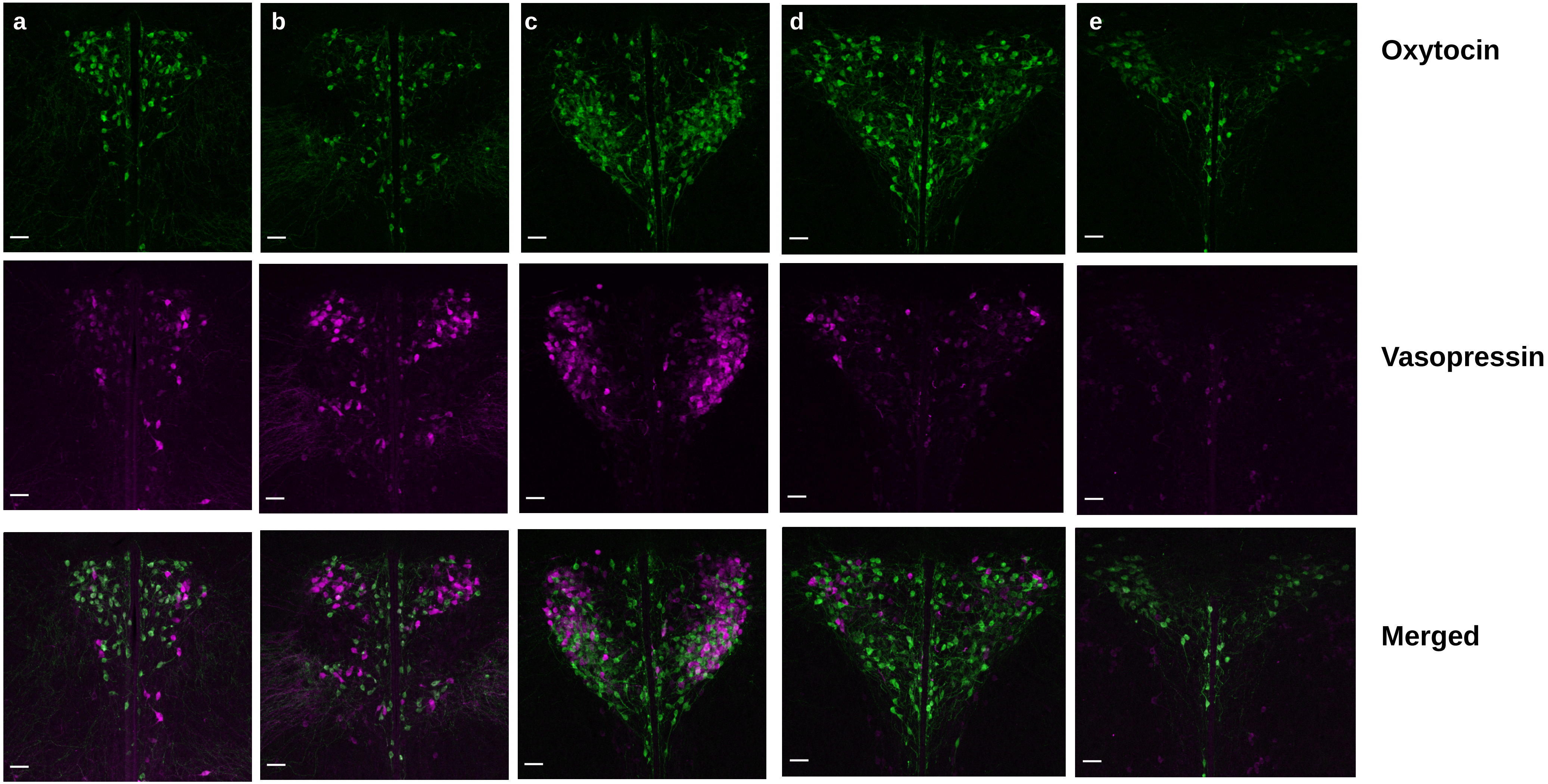


**Figure S9.** Oxytocin and vasopressin neurons' expression across the PVN in X*Y females. **a,** region 1 : anterior. **b,c**, regions 2 and 3 : intermediate. **d,e,** regions 4 and 5 : posterior.

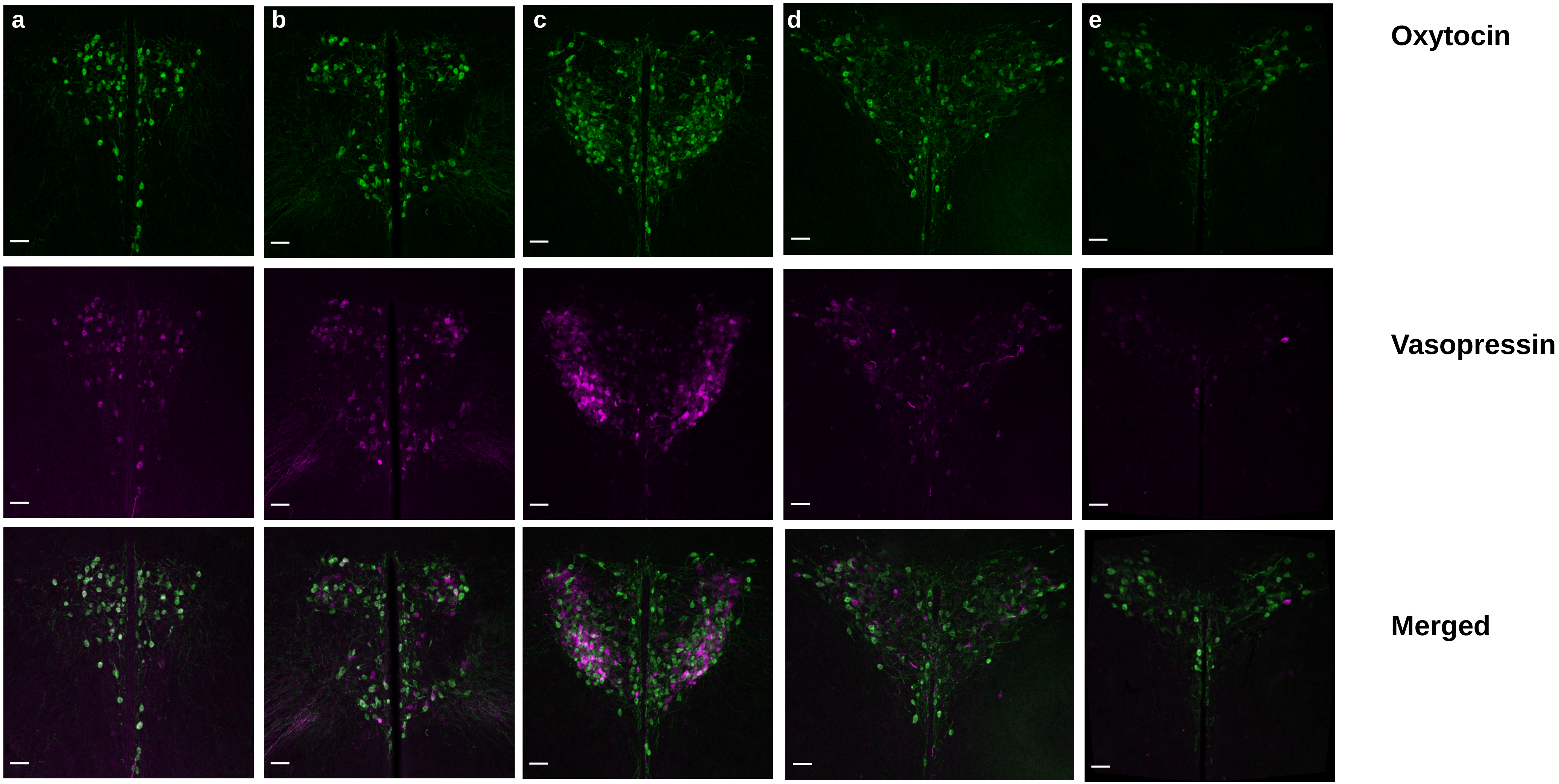


**Figure S10.** Oxytocin and vasopressin neurons' expression across the PVN in XY males. **a,** region 1 : anterior. **b,c**, regions 2 and 3 : intermediate. **d,e,** regions 4 and 5 : posterior.

| | Latency to visit (s) | | Retrieval probability | |
|---|---|---|---|---|
| | *Estimates* | *HPD (89%)* | *Estimates* | *HPD (89%)* |
| XX | 156.5 | 94.5 –223.3 | 0.47 | 0.32 –0.63 |
| XX* | 112 | 71.5– 158.6 | 0.55 | 0.42 –0.70 |
| X*Y | 51.7 | 33.6– 73.8 | 0.87 | 0.77 –0.95 |
| **Contrasts** | *ratio* | *HPD (89%)* | *Odd ratio* | *HPD (89%)* |
| XX / X*Y | 2.97 | 1.57-4.65 | 0.14 | 0.03–0.30 |
| XX / XX* | 1.4 | 0.7-2.1 | 0.74 | 0.23–1.4 |
| XX* / X*Y | 2.1 | 1.09-3.31 | 0.19 | 0.04-0.4 |

**Table S1.** Median estimates and median-based contrast estimates for pup retrieval experiments. Contrast estimates with 89% HPD including 1 are considered as not significantly different. An odd-ratio above one (or near 0) indicates a higher (or lower) behavioural likelihood in the first group of the contrast, an odd-ratio around 1 indicates no group-effect. Probabilities and latencies (s) are retrieved using a back-transformation of the model parameters. $n_{XX}$=31 , $n_{XX*}$=35, $n_{X*Y}$=35.

| Genotype | *N* | Pup retrieval<br>Visit response (%) | Retrieval response (%) | Mother-pup interactions<br>N | Carrying (%) | Grooming (%) | Crouching (%) | Huddling (%) |
|---|---|---|---|---|---|---|---|---|
| XX | 31 | 20 (64.5) | 15 (48.4) | 25 | 19 (0.76) | 20 (0.8) | 23 (0.92) | 18 (0.72) |
| XX* | 35 | 29 (82.9) | 19 (54.3) | 25 | 17 (0.68) | 22 (0.88) | 22 (0.88) | 21 (0.84) |
| X*Y | 35 | 31 (88.6) | 30 (85.7) | 25 | 17 (0.68) | 18 (0.72) | 20 (0.8) | 22 (0.88) |

**Table S2.** Group sample sizes and proportions of individuals showing a response for pup retrieval and mother-pup interactions experiments.

| | Grooming Duration (s) | | Crouching duration (s) | | Huddling duration (s) | | Pup displacement probability | |
|---|---|---|---|---|---|---|---|---|
| | *Estimates* | *HPD (89%)* | *Estimates* | *HPD (89%)* | *Estimates* | *HPD (89%)* | *Estimates* | *HPD (89%)* |
| XX | 4.87 | 2.49-7.46 | 10.98 | 5.32-19 | 4.95 | 2.15-8 | 0.76 | 0.53-0.94 |
| XX* | 5.79 | 2.89-8.70 | 15.61 | 7.73-26.5 | 7.30 | 3.43-11.5 | 0.67 | 0.42-0.89 |
| X*Y | 3.18 | 1.36-4.99 | 7.89 | 3.34-13.2 | 7.33 | 3.63-11.7 | 0.68 | 0.43-0.9 |
| **Contrasts** | *ratio* | *HPD (89%)* | *ratio* | *HPD (89%)* | *ratio* | *HPD (89%)* | *Odd ratio* | *HPD (89%)* |
| XX / X*Y | 1.42 | 0.75-2.21 | 1.36 | 0.52-2.52 | 0.71 | 0.32-1.21 | 1.54 | 0.24-5.93 |
| XX / XX* | 0.87 | 0.44-1.33 | 0.71 | 0.27–1.34 | 0.71 | 0.32-1.2 | 1.58 | 0.25-6.37 |
| XX*/ X*Y | 1.64 | 0.86-2.57 | 1.88 | 0.66-3.43 | 1 | 0.42-1.62 | 0.96 | 0.12-3.47 |

**Table S3.** Median estimates and median-based contrast estimates for mother-pup interaction investigations Contrast estimates with 89% HPD including 1 are considered as not significantly different. An odd-ratio above one (or near 0) indicates a higher (or lower) behavioural likelihood in the first group of the contrast, an odd-ratio around 1 indicates no group-effect. Probabilities and durations (s) are retrieved using a back-transformation of the model parameters. n=25 per genotype.
